# An actin-based protrusion originating from a podosome-enriched region initiates macrophage fusion

**DOI:** 10.1101/538314

**Authors:** James J. Faust, Arnat Balabiyev, John M. Heddleston, Nataly P. Podolnikova, D. Page Baluch, Teng-Leong Chew, Tatiana P. Ugarova

## Abstract

Macrophage fusion resulting in the formation of multinucleated giant cells occurs in a variety of chronic inflammatory diseases, yet the mechanism responsible for initiating macrophage fusion is unknown. Here, we used live cell imaging to show that actin-based protrusions at the leading edge initiate macrophage fusion. Phase contrast video microscopy demonstrated that in the majority of events, short protrusions (3 ± 1 μm) between two closely apposed cells initiated fusion, but occasionally we observed long protrusions (16 ± 7 μm). Using macrophages isolated from LifeAct mice and imaging with lattice light sheet microscopy, we further found that fusion-competent actin-based protrusions formed at sites enriched in podosomes. Inducing fusion in mixed populations of GFP- and mRFP-LifeAct macrophages showed rapid spatial overlap between GFP and RFP signal at the site of fusion. Cytochalasin B strongly reduced fusion and when rare fusion events occurred, protrusions were not observed. Fusion of macrophages deficient in Wiskott-Aldrich syndrome protein and Cdc42, key molecules involved in the formation of actin-based protrusions and podosomes, was also impaired both *in vitro* and *in vivo*. Finally, inhibiting the activity of the Arp2/3 complex decreased fusion and podosome formation. Together these data indicate that an actin-based protrusion formed at the leading edge initiates macrophage fusion.

## Introduction

Cell to cell fusion is an essential event in several biological processes such as fertilization, embryonic development, skeletal muscle and placenta formation, bone remodeling and stem cell differentiation (Aguilar et al., 2013; Podbilewicz, 2014). Furthermore, cell-cell fusion has been observed in a number of pathological conditions. In particular, macrophage fusion resulting in the formation of multinucleated giant cells (MGCs) is associated with numerous chronic inflammatory diseases including granulomatous infection, the foreign body reaction to implanted biomaterials, atherosclerosis, amyotrophic lateral sclerosis, cancer and others (Helming and Gordon, 2007; Anderson et al., 2008). MGCs are formed from blood monocytes recruited from the circulation to sites of inflammation where they differentiate into macrophages that undergo fusion as inflammation progresses to the chronic state. The T-helper 2 cytokine interleukin-4 (IL-4) promotes macrophages fusion *in vivo* (Kao et al., 1995) and when applied in cell culture can be used to study this process (McInnes and Rennick, 1988; McNally and Anderson, 1995). Although this *in vitro* cell system has proven invaluable to our understanding of the molecular mediators that orchestrate macrophage fusion (McNally and Anderson, 2002; Jay et al., 2007; Helming and Gordon, 2009; Milde et al., 2015), there is little information regarding the morphological changes that macrophages undergo to initiate fusion as well as the cellular mechanisms that govern this process.

MGC formation is thought to be a multistage process involving adhesion of cells to the substrate, the induction of a fusion-competent state, cellular motility, cell-cell interaction, cytoskeletal rearrangements and subsequent membrane fusion (Helming and Gordon, 2009). Most, if not all, of the steps involved in macrophage fusion appear to rely on contractile networks formed by the actin cytoskeleton. It has been shown that the fungal toxins cytochalasin B and D, which both prevent actin polymerization inhibit MGC formation in a concentration-dependent manner (DeFife et al., 1999). The importance of the actin cytoskeleton has been further corroborated by studies indicating that IL-4 activated the Rac-1 signaling pathway (Jay et al., 2007). Rac-1 is known to reorganize actin networks resulting in formation of membrane ruffles and extension of lamellipodia. Abrogation of Rac-1 activation by chemical and genetic approaches inhibited lamellipodia formation and attenuated MGC formation (Jay et al., 2007).

Several types of plasma membrane protrusions, including lamellipodia, filopodia and invadosomes can form and coexist at the leading edge of migrating cells (for review see Ridley, 2011). The formation of these protrusions is a result of actin polymerization mediated by actin nucleation-promoting factors. The primary mediators of actin polymerization that induce formation of branched networks in lamellipodia are the members of the WASp (Wiscott-Aldrich syndrome protein) family that activate the Arp2/3 complex (for review see Takenawa and Suetsugu, 2007). The Arp2/3 complex and WASp have also been implicated in filopodia formation (Takenawa and Suetsugu, 2007; Lee et al., 2010; Yang and Svitkina, 2011). Recent studies in myoblasts, cells that undergo fusion in arthropods and vertebrates have revealed many proteins that participate in Arp2/3-mediated actin polymerization and are required for fusion (Chen, 2011; Aguilar et al., 2013). In these cells, Arp2/3-mediated actin polymerization is responsible for the formation of F-actin enriched structures protruding from one cell into another cell at the site of fusion (Sens et al., 2010; Haralalka et al., 2011; Shilagardi et al., 2013). The size and the molecular composition of these protrusions (Sens et al., 2010; Chen, 2011) clearly distinguish them from filopodia and lamellipodia (Mattila and Lappalainen, 2008). Based on the presence of an actin core with a surrounding ring of adhesive proteins and their protrusive nature, the protrusions in fusing myoblasts have been called “podosome-like structures” (Onel and Renkawitz-Pohl, 2009; Sens et al., 2010). Podosomes and related structures invadopodia, collectively known as invadosomes, are ventral protrusions that form contacts with the extracellular matrix that have been identified in a variety of cell types (Linder et al., 2011; Murphy and Courtneidge, 2011). Podosomes are especially prominent in cells of the monocytic lineage, including macrophages and dendritic cells, where they have been associated with cell adhesion, migration and matrix degradation. A defining feature of podosomes is a core of actin filaments nucleated by Arp2/3 complex (Linder et al., 2000; Kaverina et al., 2003) surrounded by adhesive plaque proteins such as talin, vinculin, integrins and others (Zambonin-Zallone et al., 1989; Pfaff and Jurdic, 2001). Despite a requirement for actin polymerization in macrophage fusion, little is known about the role of actin-based protrusions during macrophage fusion.

In the present study, we reveal the existence of an actin-based protrusion that initiates IL-4-mediated macrophage fusion. Phase contrast video microscopy demonstrated that short phase dense protrusions originating at the leading edge initiated ~90% of the fusion events with the remaining events having been initiated by long protrusions. Using macrophages isolated from LifeAct mice and imaging with lattice light sheet microscopy (LLSM), we observed short actin-based protrusions originating from regions enriched in podosomes prior to macrophage fusion. Inducing fusion in mixed populations of GFP- and mRFP-LifeAct macrophages showed rapid spatial overlap between GFP/RFP signal and reorganization of the peripheral actin network at the site of fusion. Inhibiting actin polymerization with cytochalasin B impaired fusion, and when rare fusion events occurred, we observed no protrusions. Furthermore, Cdc42- and WASp-deficient macrophages fused at very low rates both *in vitro* and *in vivo* and video analysis of fusion with these cells showed no clear evidence of protrusions. Finally, inhibiting the Arp2/3 complex not only reduced fusion, but rare fusion events did not appear to be dependent on protrusions.

## Results

### Phase-dense protrusions at the leading edge precede macrophage fusion

We recently developed optical-quality glass surfaces that enabled the first time resolved views of IL-4-induced macrophage fusion and MGC formation (Faust et al., 2017; Faust et al., 2018). Using these surfaces, we showed that macrophage fusion occurred between the intercellular margins of macrophages. Furthermore, we observed a founder population of mononuclear macrophages that initiates fusion with neighboring mononuclear macrophages (Type 1 fusion). Early multinucleated cells then fuse with neighboring mononuclear macrophages (Type 2 fusion) and, finally, MGCs fuse with surrounding MGCs to form syncytia. However, due to the low-magnification views required to visualize the formation of large MGCs, the mechanism underlying this process remained obscure.

To visualize structures between the intercellular margins of fusing macrophages in detail, we initially used phase-contrast video microscopy with intermediate magnification objectives. In this series of experiments, we used primary macrophages isolated from the inflamed mouse peritoneum (Helming and Gordon, 2007a; Podolnikova et al., 2016) in order to avoid robust cell division observed in cultures of macrophage cell lines. Analyses of Type 1 fusion events (n=31) revealed the existence of phase-dense protrusions immediately preceding macrophage fusion (Figure 1A). For the majority of events (~90%), short protrusions (3 ± 1 μm) initiated fusion (Figure 1A and Video 1). However, occasionally we observed long protrusions (16 ± 7 μm) (Figure 1B and Video 2). Furthermore, several patterns of macrophage fusion emerged. Among Type 1 events, fusion most frequently (34%) occurred between the leading edge of one cell and the cell body of another followed by fusion from the leading edge to the lagging end (23%) and between the leading edges of two closely apposed cells (20%) (Table 1 and Videos 3-5). Similar patterns of fusion were observed in the Type 2 group (n=39) (Table 1). The other patterns included fusion between the cell-cell bodies and fusion between lagging ends, among others. The time required from first intercellular contact until full nuclear integration between two macrophages was also similar for Type 1 and 2 fusion events (52 ± 26 min vs. 57 ± 22 min, respectively). We also characterized fusion of macrophages using Permanox™ (n=48), a permissive plastic surface that is widely used to induce macrophage fusion (Helming and Gordon, 2007a; Milde et al., 2015). The only significant difference observed between macrophages undergoing fusion on optical-quality glass and Permanox™ plastic was the total average time as determined by first observable contact between macrophages to full integration, when Type 1 and Type 2 events were pooled (54.5 ± 24 min vs. 64.3 ± 23 min, respectively).

**Table 1.**
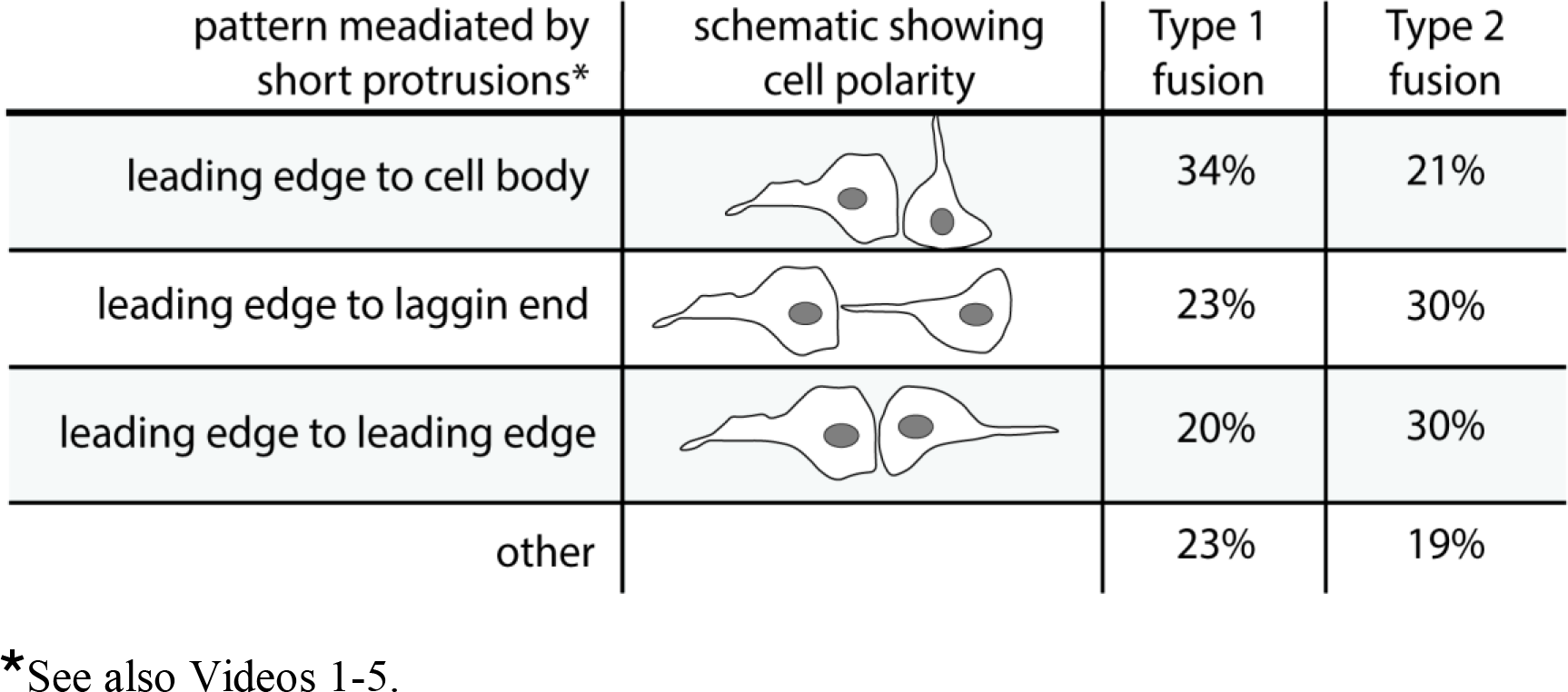
Patterns of macrophage fusion

| pattern mediated by short protrusions* | schematic showing cell polarity | Type 1 fusion | Type 2 fusion |
| --- | --- | --- | --- |
| leading edge to cell body |  | 34% | 21% |
| leading edge to lagging end |  | 23% | 30% |
| leading edge to leading edge |  | 20% | 30% |
| other |  | 23% | 19% |

\*See also Videos 1-5.

**Figure 1:**
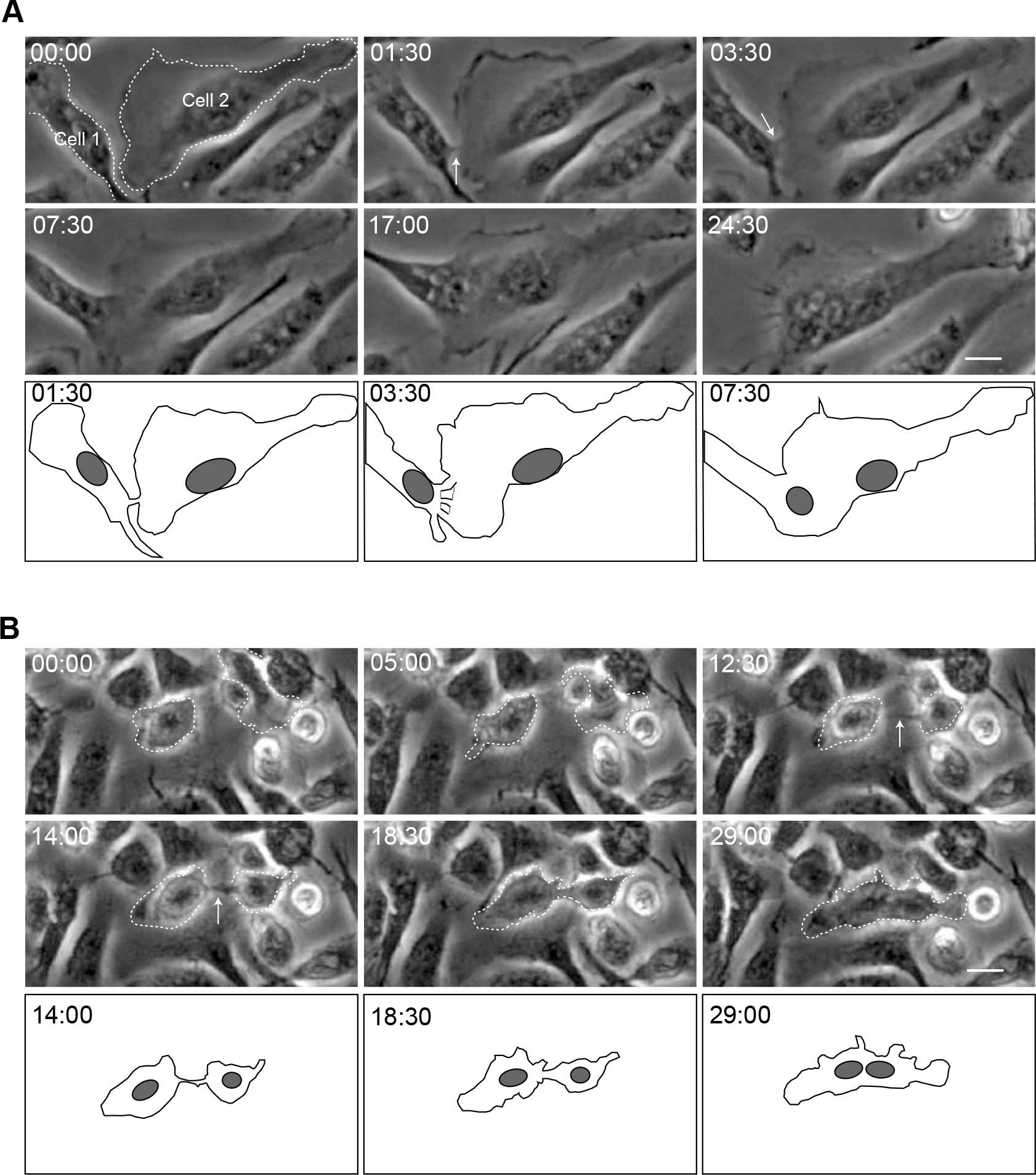
Phase-dense protrusions initiate macrophage fusion. **(A)** Live imaging of mononuclear macrophages undergoing fusion. Macrophages were isolated from the mouse peritoneum 3 days after TG injection, plated on a 35-mm Fluorodish and fusion was induced by IL-4. Mononuclear macrophage (Cell 1) extends a short phase-dense protrusion (white arrow) toward another mononuclear macrophage (Cell 2) immediately before fusion. The lower panel is a diagram of frames at 1:30, 3:30 and 7:30 min illustrating morphological aspects of the fusion process. See also Video 1. **(B)** Macrophages undergoing Type 1 fusion can occasionally extend a long protrusion (white arrow) to initiate fusion. The lower panels show diagrams of frames at 14:00, 18:30 and 29:00 min. In each micrograph, time is shown in minutes:seconds. The scale bars are 10 μm. See also Video 2.

### An actin-based protrusion initiates macrophage fusion

Membrane protrusions at the leading edge are actin-based structures. Furthermore, F-actin is known to be important for the formation of MGCs (DeFife et al., 1999; Jay et al., 2007). To determine if the protrusions we observed in phase-contrast micrographs were actin-based structures, we examined macrophages isolated from the inflamed peritoneum of LifeAct mice using LLSM (Chen et al., 2014). We first confirmed that LifeAct faithfully reported the distribution of F-actin in macrophages by comparing the distribution of eGFP-LifeAct and Alexa 568-conjugated phalloidin (Supplemental Figure 1). Using LLSM, we reveal waves of eGFP-LifeAct puncta emanating from the center of the cell to the cell periphery prior to fusion (Video 6). At the time of apparent fusion, one wave of LifeAct puncta advanced into a neighboring cell (Figure 2A and Video 6). In fixed specimens, actin puncta of similar dimensions at cellular margins contained rings of talin and vinculin circumscribing a central actin core in both mononuclear macrophages and MGCs (Supplemental Figure 2; vinculin shown), suggesting that structures we observed in living cells were podosomes. When we analyzed the site of fusion (Figure 2B; 29:11-29:24 min; the boxed areas in Figure 2A) with the maximum temporal resolution achievable with LLSM under the conditions of our experiments (~1.5 s per image for several hours), we observed a finger-like enrichment of LifeAct that extended into the neighboring cell during fusion (Figure 2B; arrows). This protrusive structure fanned from the initial point of contact as fusion proceeded (Figure 2B, 29:24 min).

**Figure 2:**
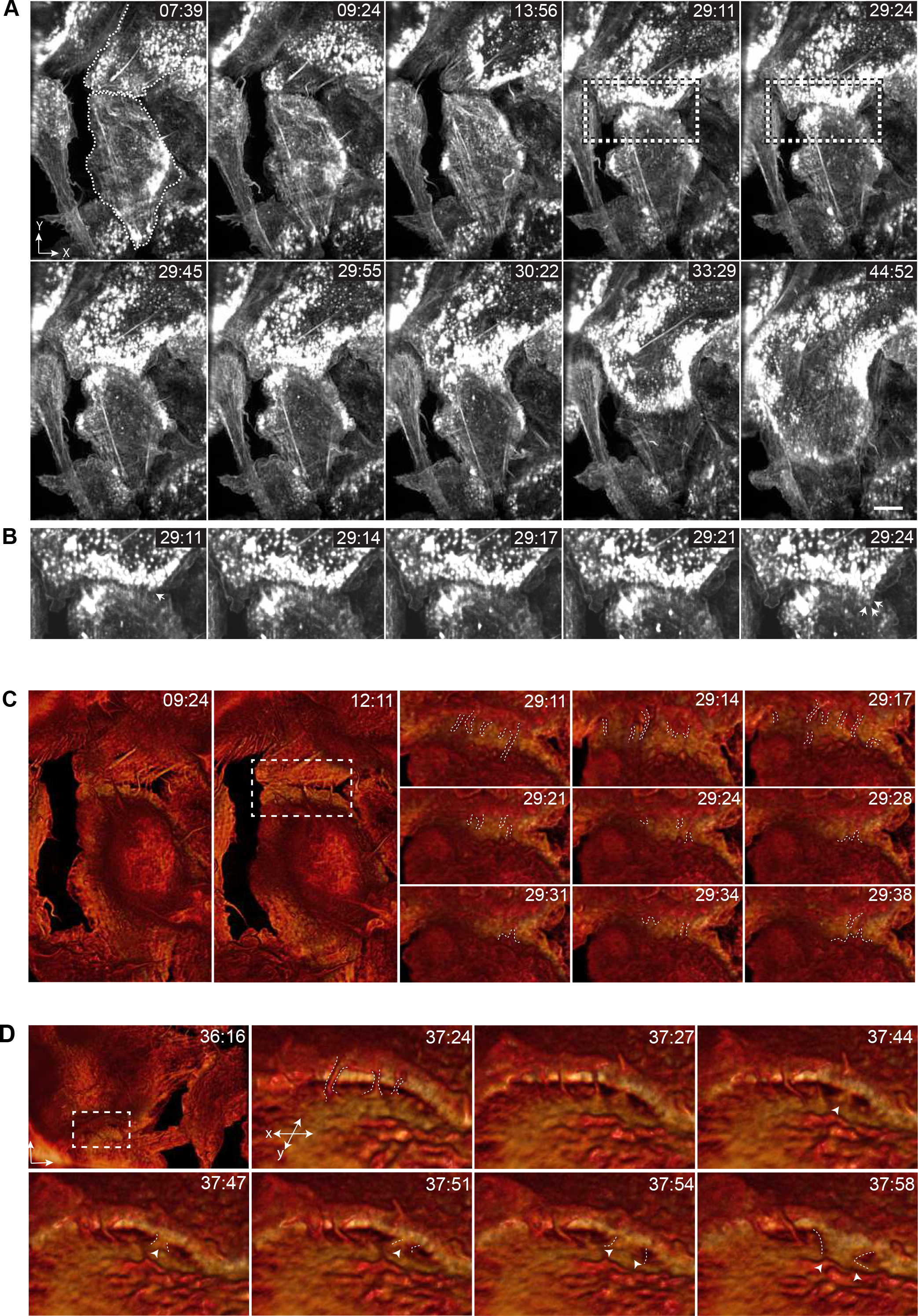
An actin-based protrusion precedes macrophage fusion. **(A)** Lattice light sheet microscopy of IL-4-induced fusion of macrophages expressing eGFP-LifeAct. See also Video 6. **(B)** Enlarged images of events occurring between 29:11 and 29:24 min (boxed regions in A). All images shown are maximum-intensity projections. The scale bar in A is 5 μm. **(C)** *En face* isosurface renders of LLSM data from A. The boxed region at 12:11 min corresponds to the subsequent micrographs showing the fusion progress (29:11-29:38 min). Note numerous protrusions (outlined by white dashes) formed between apposing cells. See also Video 7. **(D)** Surface renders of LLSM data from another area of fusing macrophages showing contact of a protrusion (white arrow; 37:27-37:51 min) followed by its apparent expansion (white arrows; 37:54-37:58). Time in each micrograph is shown as minutes:seconds. See also Video 8.

To visualize the dynamics of the actin cytoskeleton prior to fusion and observe the interface between fusing cells, we used a maximum intensity isosurface render of eGPF-LifeAct applied to the area shown in Figure 2A (Figure 2C and Video 7). We observed numerous thin protrusions between two interacting cells after initial contact (Figure 2C; 9:24 and 12:11 min). The protrusions appeared to contact apposing cells by rounds of extension and retraction, which continued until seconds before the cells fused (Figure 2C; shown for 29:11-29:38 min; individual protrusions are outlined). Close apposition of these cells in diffraction-limited space precluded visualizing the role of protrusions during the fusion process. Analysis of another area also revealed the presence of protrusions extending and retracting between two cells (Figure 2D; white outline; and Video 8). In this case, however, separation between the cells made it possible to observe an actin-based protrusion that initiated fusion (Figure 2D; 37:27-37:51 min; white arrows). Gradual expansion of the actin network that followed at this site could potentially be attributed to the local expansion of the protrusion, the formation of a fusion pore, or both (Figure 2D, 37:54-37:58 min; arrows).

### F-actin incorporation is asymmetric during macrophage fusion

To determine how actin is integrated during the fusion process, we mixed equal numbers of eGFP-LifeAct and mRFP-LifeAct macrophages, induced fusion, and imaged the process with LLSM (Video 9). When we visualized cells in this mixing assay at early time points, we observed no overlap of GFP and RFP emission in diffraction limited space (Figure 3A; 5:57 min; two adjacent mononuclear cells in the right bottom quadrant are outlined). However, at the time of apparent fusion (Figure 3A; 06:14 min), we observed overlap of GFP and RFP in overlays at the cell margins, which became more apparent as fusion proceeded (Figure 3A; 6:31-7:22 min). Further, we observed reorganization of actin in cells undergoing fusion that first clearly appeared at 09:36 min and was completed by 20:09 min, suggesting mixing of the cytoplasm (Figure 3A).

**Figure 3:**
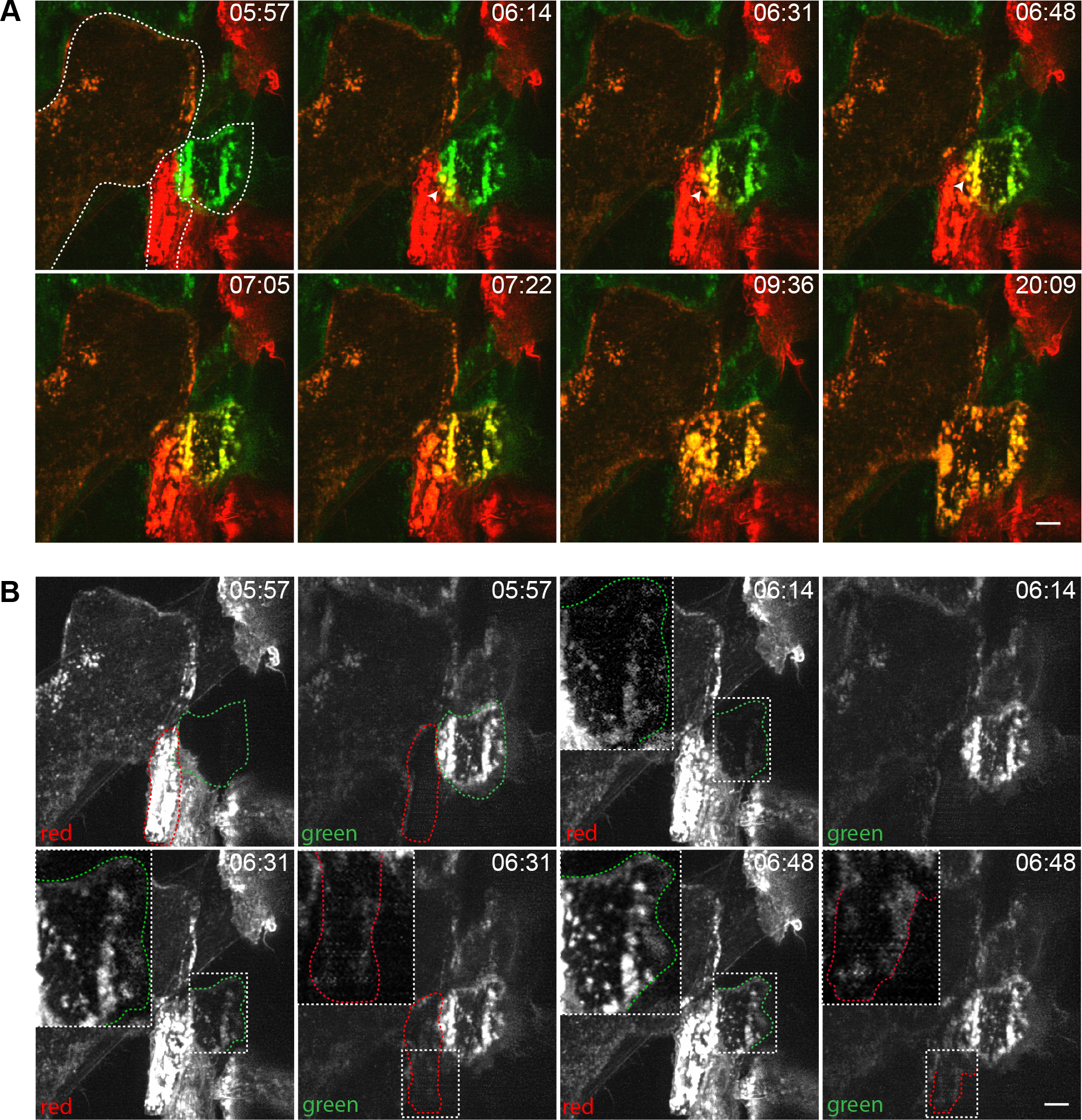
Mixing macrophages expressing eGFP-LifeAct and mRFP-LifeAct shows asymmetric actin integration. **(A)** Macrophage fusion in a mixed population of eGFP/mRFP-LifeAct macrophages. Two mononuclear cells expressing eGFP-LifeAct and mRFP-LifeAct undergoing fusion are outlined in the image taken at 5:57 min. Sites of eGFP- and mRFP-LifeAct integration are indicated by white arrows at 6:14-6:48 min. **(B)** Split-channel view of the fusion event shown in A. The boxes outlined in white dashed lines in the images taken at 6:14-6:48 min correspond to the areas where integration of mRFP-LifeAct into the eGFP-LifeAct-containing cells is observed. The enlarged images of the same areas are shown as insets. The scale bars are 5 μm. See also Videos 9 and 10.

Separating GFP and RFP emission and analyzing fusion in the two cells outlined in green and red revealed an asymmetry in the fusion process (Figure 3B and Video 10). Prior to fusion (05:57 min), we observed no GFP signal in the cell outlined in red and vice versa. However, at 06:14 min, signal from RFP appeared in the green outline traversing the entire length of the cell (Figure 3B; magnified *inset* shows the outline of the eGFP-LifeAct macrophage). At the same time, we were unable to detect green signal within the outline of the red cell. Only after an additional 17 s (06:31 min) did we begin to see low levels of GFP signal spatially overlapping boundaries of the cell outlined in red, which appeared to enrich as time progressed (06:48 min). Thus, it appears that one of the two cells more actively integrates cytoplasm than the other does.

In many cases, we found reorganization of a peripheral actin network at the intercellular interface coinciding with the formation of an opening between two cells. Shown in Figure 4A is fusion of a multinucleated cell (Cell 1) and a binucleated cell (Cell 2). In this case, before fusion (16:19 min), both cells already contained mixed eGFP-LifeAct and mRFP-LifeAct that appeared enriched at the cell periphery. At the point of apparent fusion (20:09-20:24 min), the interface that partitioned the cells was deconstructed (white arrows). Mixed LifeAct, which initially was at the interface between the cells disassembled to create an opening that gradually expanded (20:38 min and onward). Intensity scans of the leading edge interface of Cell 2 shows a drop in intensity after apparent fusion, indicating loss of the interface (Figure 4B and Video 11).

**Figure 4:**
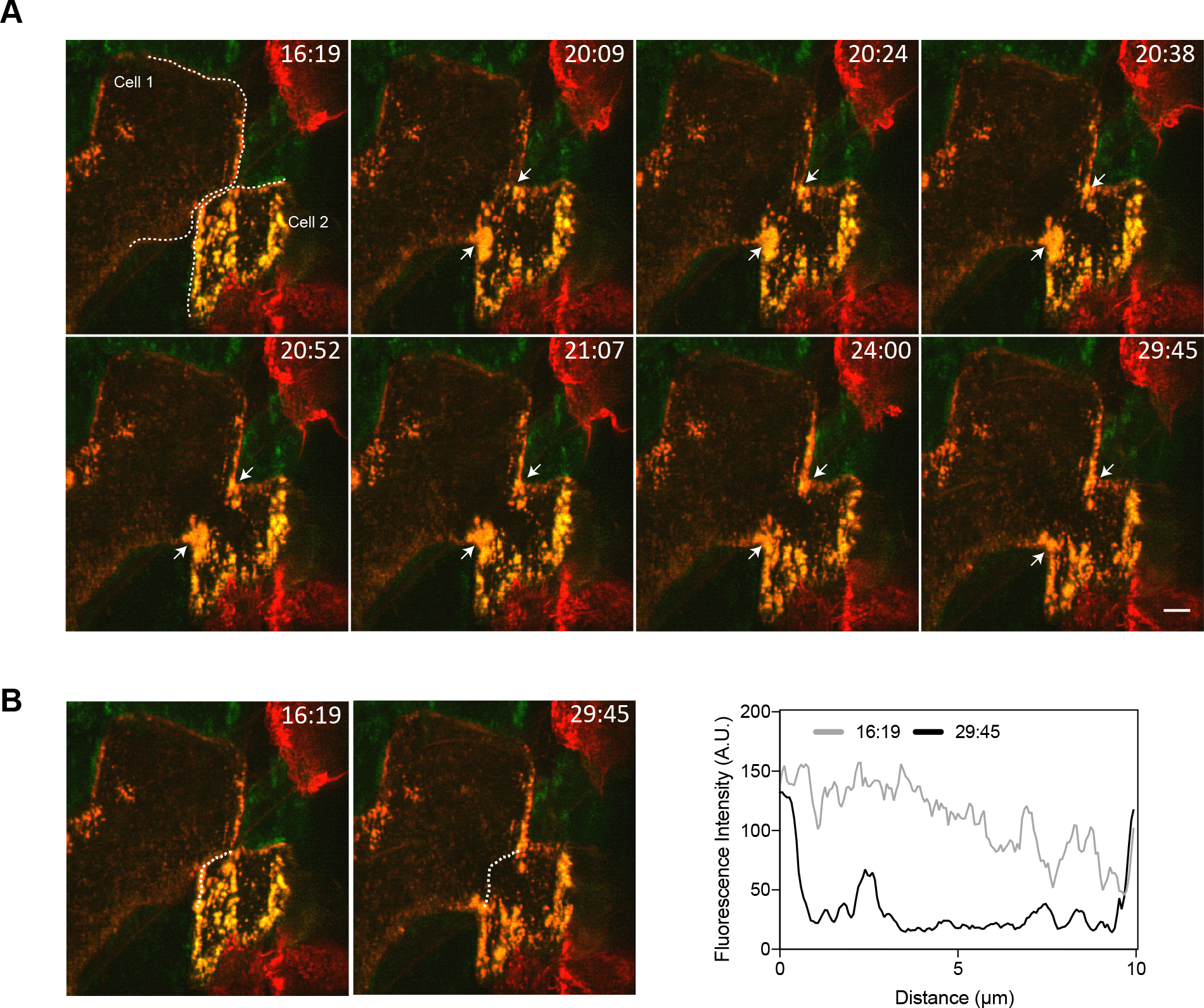
Reorganization of the peripheral LifeAct network during fusion. **(A)** Fusion was induced by IL-4 in the culture containing eGFP-LifeAct and mRFP-LifeActs expressing macrophages. Two cells (Cell 1 and Cell 2) that eventually undergo fusion are outlined in white dashes. The white arrows point to the site of fusion. As time progresses, LifeAct signal near the site of fusion appears to reorganize to expand the pore. The scale bar is 5 μm. **(B)** Intensity scans of the pore before and after apparent fusion. One pixel line scans of Cell 2 were used to generate the intensity profile at the times indicated and corresponds to the dashed lines. See also Video 11.

### Organizers of actin-based protrusions are critical for macrophage fusion *in vitro*

Since protrusions precede macrophage fusion, we sought to determine whether these actin-based structures are causal mediators of macrophage fusion. As a first step, we examined the role of F-actin, the cytoskeleton underlying membrane protrusions, in macrophage fusion. Consistent with previous data (DeFife et al., 1999), treatment of cells with cytochalasin B reduced MGC formation. Quantification of the fusion index indicated that at a concentration as low as 2.5 μM, cytochalasin B decreased fusion by ~3-fold from 29 ± 10% to 10 ± 5% for untreated vs. treated cells, respectively (Figure 5A and B). Furthermore, the majority of treated cells had a bulbous shape compared to a flattened shape of untreated cells (Figure 5A, *bottom panels*). Importantly, when we recorded fusion in the presence of cytochalasin B using phase contrast video microscopy, we were unable to observe phase-dense protrusions preceding fusion. Rather, some cells in close apposition appeared to passively undergo fusion (Figure 5C).

**Figure 5:**
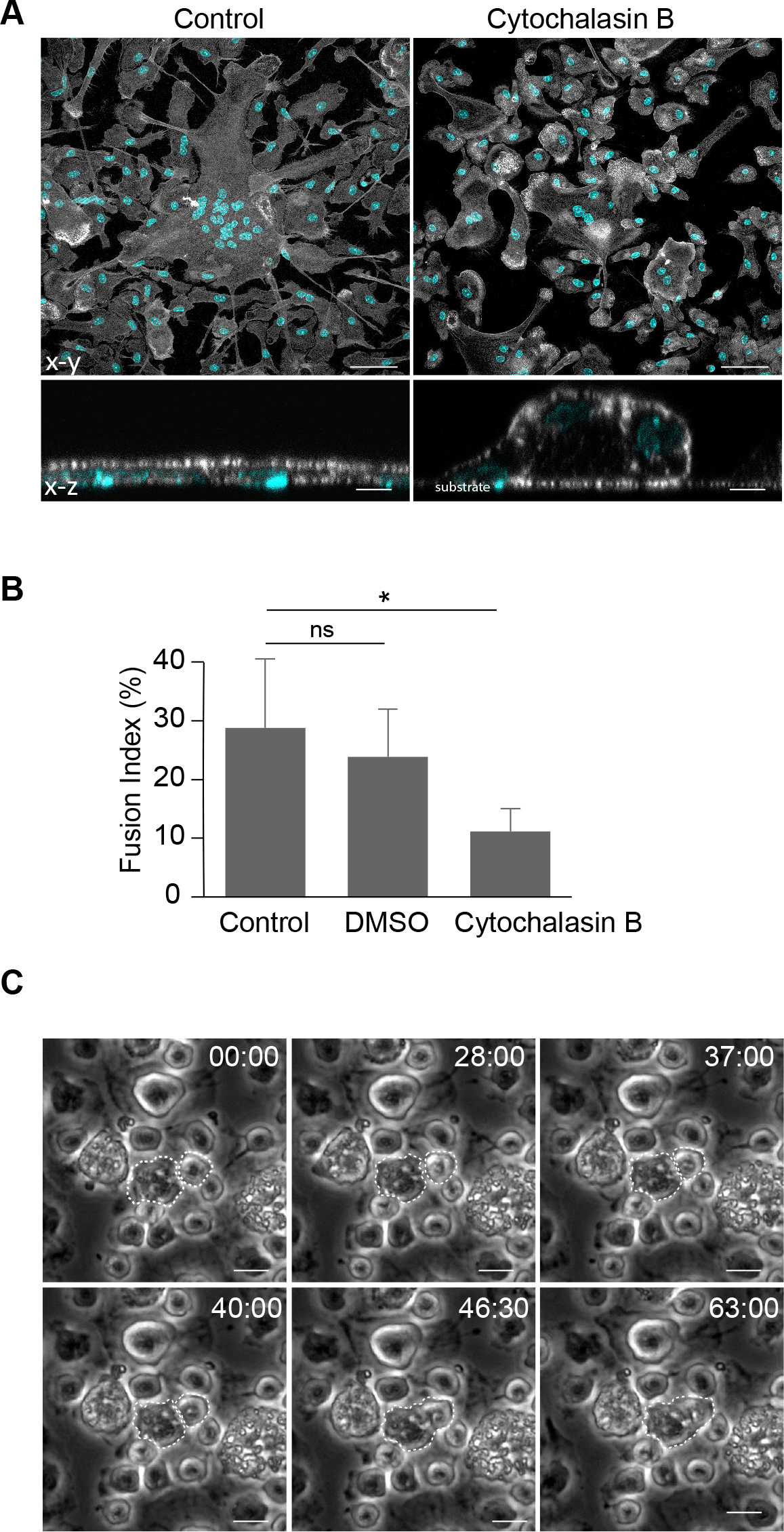
Cytochalasin B treatment reduces macrophage fusion. **(A)** Confocal micrographs of control untreated and cytochalasin B-treated (2.5 μM) macrophages 24 hours after incubation in the presence of IL-4. The cells were labeled with Alexa Fluor 488-conjugated phalloidin (white) and DAPI (teal). The bottom panels show the x-z sections of control untreated (*left*) and cytochalasin B-treated (*right*) MGCs. The scale bars are 50 μm and 10 μm in the in the upper and lower panels, respectively. **(B)** Quantification of the fusion index in the population of untreated and cytochalasin B-treated macrophages. DMSO was used as a vehicle control for cytochalasin B treatment. Results shown are mean ± SD from three independent experiments. *p<0.01. **(C)** Single cytochalasin B-treated macrophages that undergo fusion do not form protrusions. The cells that fuse are outlined. The scale bars are 10 μm.

It is well known that Cdc42 orchestrates filopodia formation. In addition, Cdc42 and PIP2 activate WASp to trigger downstream Arp2/3-mediated actin polymerization to form lamellipodia, podosomes and other protrusions (Campellone and Welch, 2010; Mattila and Lappalainen, 2008; Linder et al., 2011). If actin-based protrusions are involved in macrophage fusion, then perturbing the function of these upstream regulatory proteins should inhibit macrophage fusion. To test this prediction, we isolated macrophages from a WASp^−/−^ mouse and examined IL-4-induced macrophage fusion at various time points. Figures 6A and 6B show that fusion of WASp-deficient macrophages was strongly impaired. Compared to WT macrophages, the degree of fusion of WASp-deficient macrophages at every time point tested (24, 48 and 72 hours) was ~6-fold less. WASp deficiency also inhibited the formation of podosomes in fusing macrophages (Figure 6C). Gross morphology and the degree of macrophage adhesion after 2.5 hours in culture did not appear to be significantly different from WT macrophages (Figure 6A; t=0). Using phase contrast video microscopy performed during first 24 hours after IL-4 addition we were able to observe rare fusion events, which appear to have occurred by a protrusion-independent mechanism (Figure 6D).

**Figure 6:**
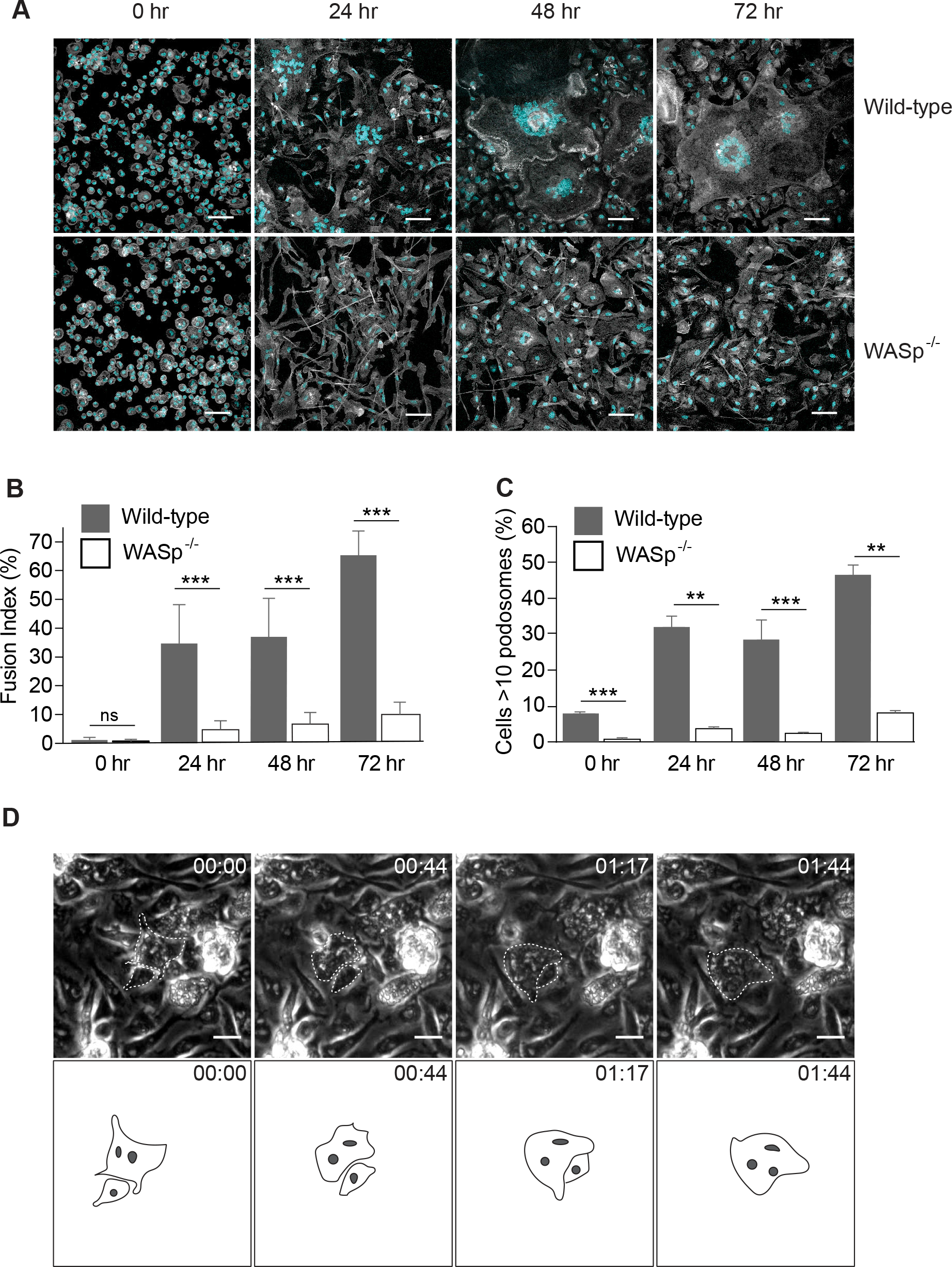
WASp is required for macrophage fusion *in vitro*. **(A)** Fusion of wild-type and WASp-deficient macrophages at various time points after the addition of IL-4. After 24, 48 and 72 hours, cells were fixed and labeled with Alexa 488-conjugated phalloidin (white) and DAPI (teal). The scale bars are 50 μm. **(B)** The time-dependent fusion indices for wild-type and WASp-deficient macrophages. Results shown are mean ± SD from three independent experiments. Three to five random 20x fields were used per sample to count nuclei. ***p<0.001. **(C)** The time-dependent podosome formation in fusing macrophages. Fraction of cells with >10 podosomes for each time point was calculated. Four random 20x fields each containing ~200-300 cells were used to count podosomes. **p<0.01, ***p<0.001. **(D)** Live imaging of IL-4-treated WASp-deficient macrophages. In each micrograph, time is shown in hours:minutes. A rare fusion event detected in the population consisting of ~1200 macrophages is shown.

We next examined whether Cdc42 is required for macrophage fusion using macrophages isolated from myeloid cell-specific Cdc42^−/−^ mice (Figure 7, A and B). As shown in Figure 7, C and D, at 72 hours, we observed a ~2-fold decrease in fusion of Cdc42-deficient macrophages compared to control Cdc42^loxP/loxP^-counterparts. Similar to WASp, Cdc42 deficiency also strongly reduced the formation of podosomes (Figure 7E).

**Figure 7:**
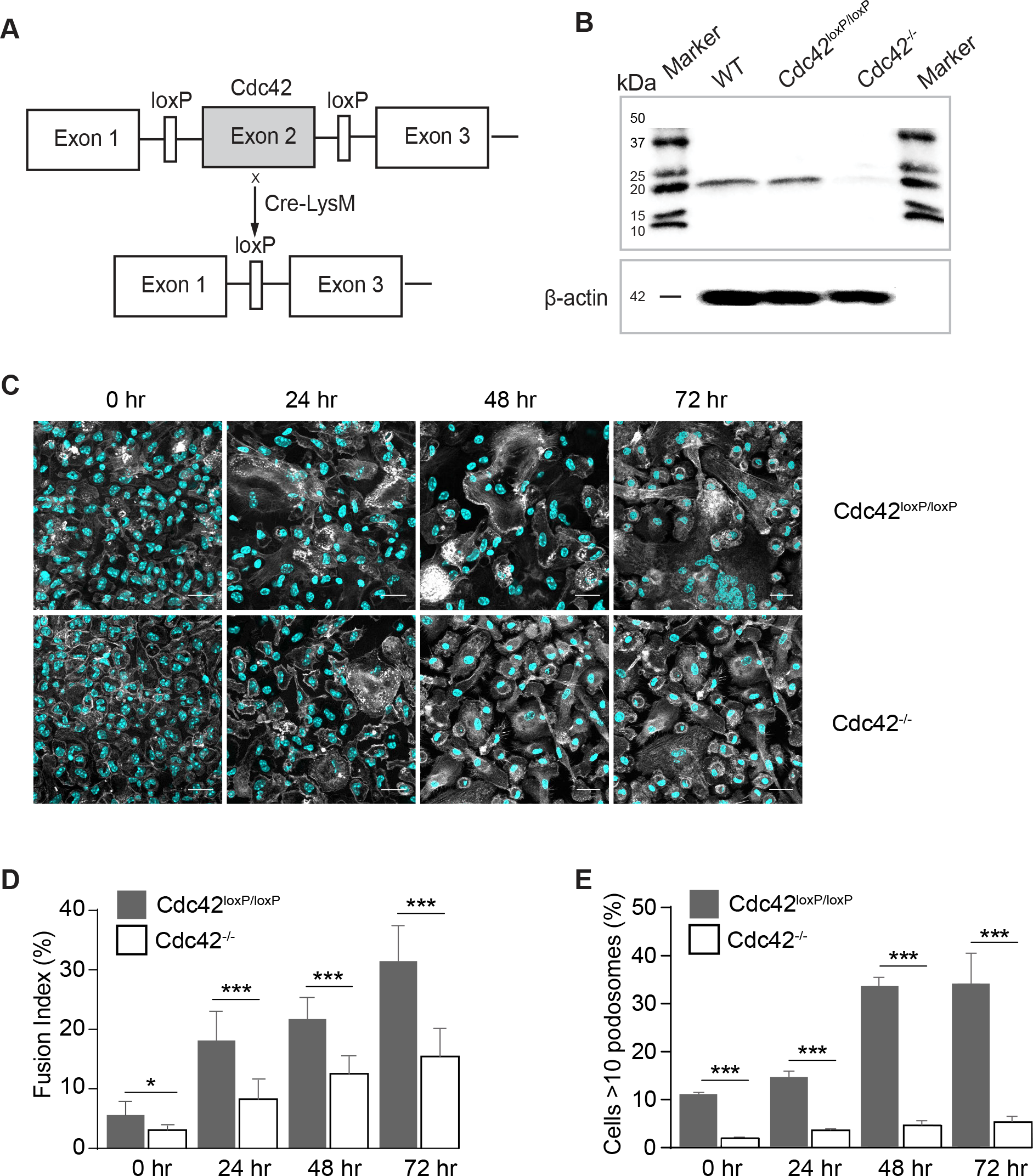
Loss of Cdc42 in macrophages results in impaired fusion *in vitro*. **(A)** Schematic diagram of the generation of a myeloid cell-specific Cdc42^−/−^ mouse. Conditional gene-targeted mice with exon 2 of *Cdc42* gene flanked by a pair of loxP sequences (Yang et al., 2007) were cross-bred with LysMcre mice to allow *Cdc42* gene excision in myeloid cells. **(B)** Cdc42 deletion in isolated macrophages was examined by SDS-PAGE (11% gel) followed by Western blotting using anti-Cdc42 rabbit monoclonal antibody (ab187643; Abcam). **(C)** Fusion of wild-type and Cdc42-deficient macrophages at various time points after the addition of IL-4. Cells were fixed after 24, 48 and 72 hours and labeled with Alexa 488-conjugated phalloidin (white) and DAPI (teal). The scale bars are 50 μm. **(D)** Time-dependent fusion indices of wild-type and Cdc42-deficient macrophages. Results shown are mean ± SD from three independent experiments. Three to five random 20x fields were used per sample to count nuclei. *p < 0.05, ***p < 0.001. **(E)** The time-dependent podosome formation in fusing macrophages. Fraction of cells with >10 podosomes for each time point was calculated. Four random 20x fields each containing ~200-300 cells were used to count podosomes. ***p < 0.001.

We next determined whether inhibition of Arp2/3 results in impaired macrophage fusion. As shown in Figure 8A, the Arp2/3 specific inhibitors CK-636 and CK-548 (Nolen et al., 2009) blocked macrophage fusion in a dose-dependent manner. The IC_50_ values for CK-636 and CK-548 inhibition were 13 ± 0.5 and 15 ± 0.7 μM, respectively, for 72-hour cultures. In addition, consistent with previous reports (Nolen et al., 2009), inhibition of Arp2/3 decreased the number of podosomes with both inhibitors exerting similar effects (IC_50_= 9.5 ± 0.5 μM for CK548 and 12 ± 0.5 μM for CK636; Figure 8B). Although ~40% fusion was observed in the presence of 15 μM CK-636, MGCs did not account for the majority of fusion events. Rather, binucleated cells were the predominant cell type, which formed by a protrusion-independent mechanism (Figure 8C).

**Figure 8:**
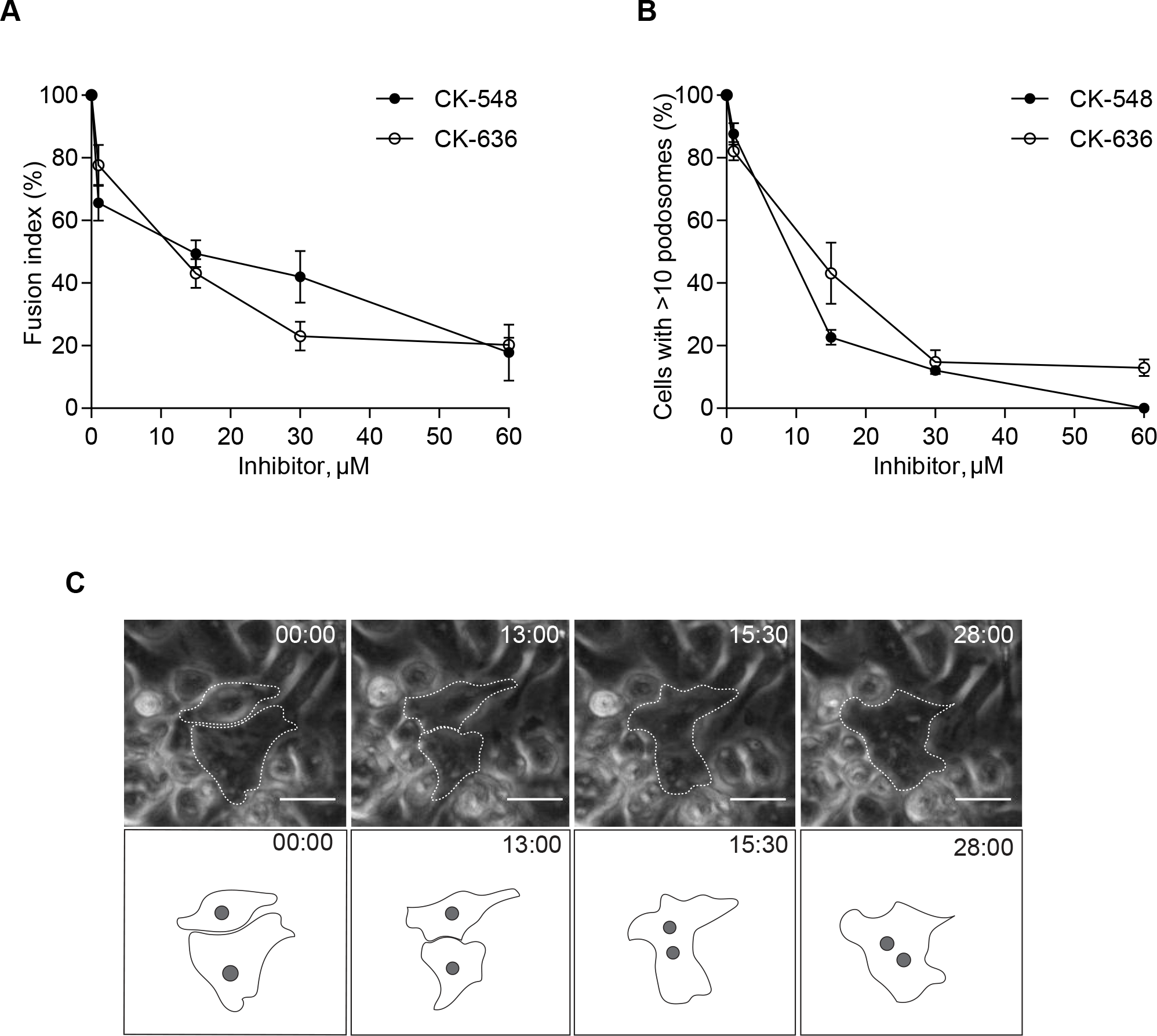
Inhibition of actin assembly by CK-636 and CK-548 decreases macrophage fusion and podosome formation. **(A)** Different concentrations of the Arp2/3 inhibitors CK-636 and CK-548 were added to macrophages at the onset of fusion induction with IL-4. Control cells were treated with DMSO. The fusion rates were determined after 72 hours from confocal images of samples labeled with Alexa 488-conjugated phalloidin and DAPI. Results shown are mean ± SD from three independent experiments. Fusion of control cells was assigned a value of 100%. Three to five random 20x fields were used per sample to count nuclei. **(B)** Effect of inhibitors (each at 15 μM) on podosome formation. Fraction of cells with >10 podosomes was calculated and normalized to DMSO control. Results shown are mean ± SD from three independent experiments with three to five random 20x fields used per sample to count cells (100-150 cells/field). **(C)** Live imaging of macrophages treated with 15 μM Arp2/3 inhibitor CK-548. In each micrograph, time is shown in minute:sec. A single fusion event detected is shown.

### WASp- and conditional Cdc42-deficient mice do not initiate a robust foreign body response to implanted materials

Further evidence for the involvement of WASp and Cdc42 in macrophage fusion was obtained in *in vivo* experiments. Macrophage fusion leading to the formation of MGCs is a hallmark of the foreign body reaction that follows the implantation of vascular grafts and other engineered devices (Anderson et al., 2008; McNally and Anderson, 2011). To confirm that WASp and Cdc42 are important for the formation of MGCs *in vivo*, we implanted polychlorotrifluoroethylene (PCTFE) into the peritoneum of WT, WASp^−/−^ and conditional Cdc42-deficient mice and retrieved the implants 14 days later. Visualization of cells covering the surface of explants revealed a large number of MGCs on materials implanted into WT, but not WASp^−/−^ mice (Figure 9, A and B). We found a ~5-fold difference in the fusion index between WT and WASp-deficient macrophages (36 ± 6% vs.7 ± 4% for WT and WASp-deficient cells, respectively) (Figure 9B). No significant difference in the number of cells recruited into the peritoneum of WT and WASp^−/−^ mice was found (Figure 9C). Conditional deletion of Cdc42 also strongly impaired multinucleation with a ~6-fold difference between Cdc42^loxP/loxP^ and Cdc42^−/−^ cells (33% ± 12 vs 5.9% ± 4.0) (Figure 9D and 9E). The number of cells in the peritoneum 14 days after implantation was not significantly different between the two strains of mice (Figure 9F). Together these data indicate that WASp and Cdc42 are essential *in vivo* for a robust foreign body reaction.

**Figure 9:**
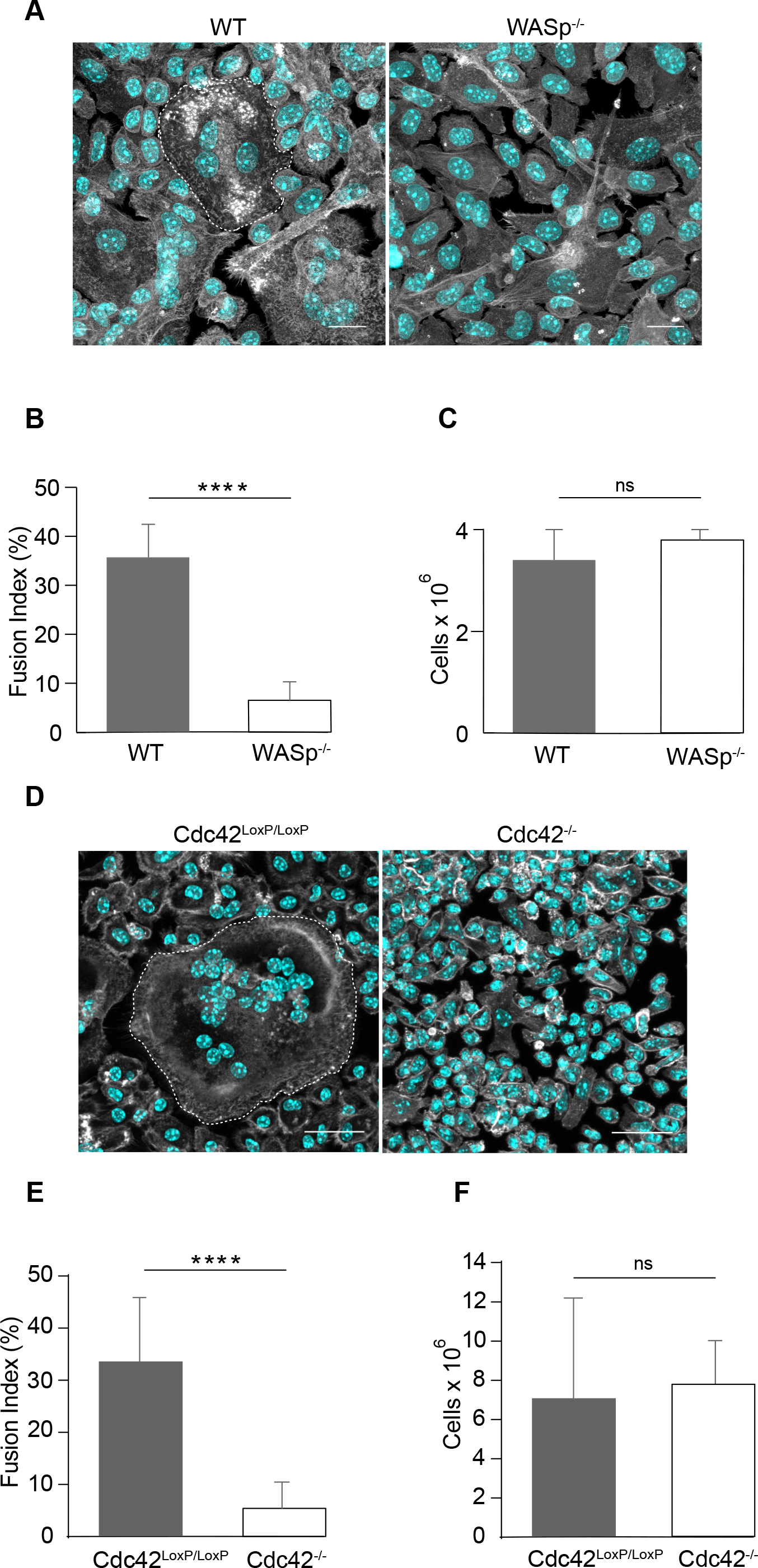
Fusion of WASp- and Cdc42-deficient macrophages is severely impaired *in vivo*. **(A)** Micrographs of macrophages on PCTFE surfaces retrieved 14 days after implantation in wild-type and WASp^−/−^ mice. The scale bars are 7.5 μm. **(B)** Quantification of the fusion index of macrophages on PCTFE surfaces retrieved from Wasp^−/−^ mice. Results shown are mean ± SD from three independent experiments. Three to five random 20x fields were used per sample to count nuclei. ****p < 0.0001. **(C)** Number of cells collected from the peritoneum of wild-type and WASp^−/−^ mice before implants were retrieved. Results shown are mean ± SD from three independent experiments. Three to five random 20x fields were used per sample to count cells from Wright stained cytospin preparations. No significant difference was observed. **(D)** Micrographs of macrophages on PCTFE surfaces retrieved 14 days after implantation in wild-type mice and mice with Cdc42-deficiency in myeloid cells. The scale bars are 30 μm. **(E)** Quantification of the fusion index of macrophages on surfaces implanted into Cdc42^−/−^ mice. Results shown are mean ± SD from three independent experiments. Three to five random 20x fields were used per sample to count nuclei. ****p< 0.0001. **(F)** Number of cells collected from the peritoneum of wild-type and Cdc42^−/−^ mice before implants were retrieved. Results shown are mean ± SD from three independent experiments. Three to five random 20x fields were used per sample to count cells. No significant difference was observed.

## Discussion

Despite the long history of research on MGCs highlighted by the fact that these cells are often observed in many diseases, the molecular and cellular mechanisms of macrophage fusion remain poorly understood. While previous studies focused mainly on the identification of fusion effector molecules in macrophages (Helming and Gordon, 2009; Vignery, 2011) little effort has been expended to elucidate the involvement of the actin cytoskeleton, which has been shown to be a driving force in other types of cells undergoing fusion (Aguilar et al., 2013; Podbilewicz, 2014). Here, we used live cell imaging of macrophages to visualize actin-based structures formed at the site of contact between two fusing cells. We show for the first time that IL-4-induced macrophage fusion is initiated by a single protrusion that more often (~80% events) extends from the leading edge of one cell in close contact with another cell. The majority of fusion-competent protrusions are short although long structures were also observed.

Fusion-competent protrusions appear to originate from cells that become enriched in peripheral podosomes. In these cells, podosomes emanate from the interior and move in a wave-like manner to the periphery where they accumulate at the site destined for fusion (Figures 2, A and B and Video 6). Before the arrival of podosomes, the interface between two cells is filled with thin protrusions that undergo rounds of extension and retraction (Figures 2C and 2D; Video 7; and schematically shown in Figure 10A). This activity continues for some time until podosomes in the cell that initiates fusion arrive and align along the plasma membrane (schematically shown in Figure 10B). Shortly after, one of the protrusions initiates fusion (Figure 10, C and D).

**Figure 10:**
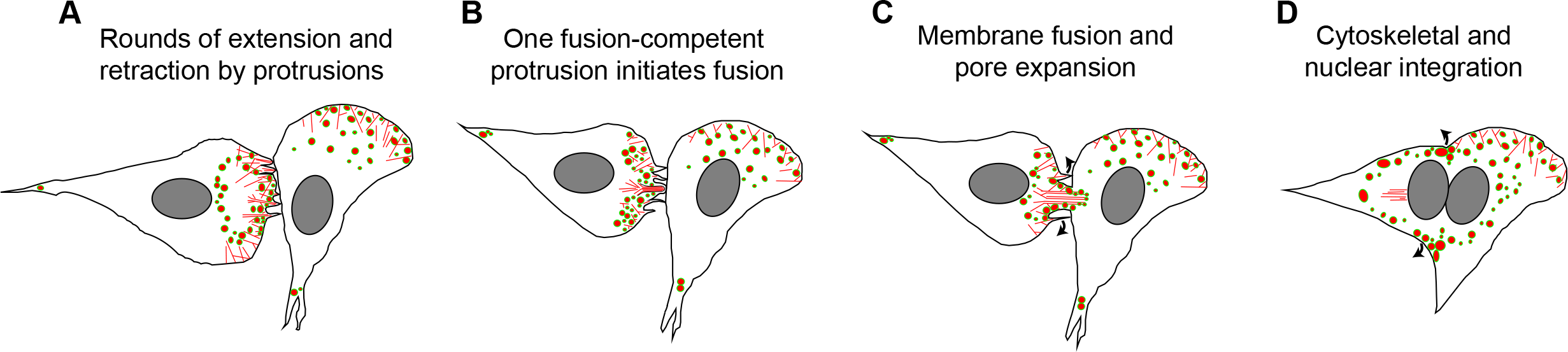
Model for macrophage fusion initiated by actin-based protrusions. **(A)** The interface between two closely apposing macrophages contains numerous thin actin-based protrusions. A wave of podosomes moves from the interior of the cell that initiates fusion to the site of cell-cell contact. **(B)** Once podosomes arrive and align at the site of contact with the acceptor macrophage, membrane fusion is initiated by a single protrusion, and podosomes from the donor macrophage advance into the acceptor macrophage. **(C)** The actin network at the site of fusion rapidly reorganizes to expand the initial pore. **(D)** The actin cytoskeleton and nuclei undergo directional integration.

Recently, a number of studies have reported the involvement of membrane extensions during osteoclastogenesis (Oikawa et al., 2012; Soe et al., 2015; Wang et al., 2015). Using RAW264.7 macrophages differentiated into osteoclasts by M-CSF and RANKL, Oikawa et al demonstrated that cell-cell fusion was mediated by short membrane protrusions that emerged from the sealing belt, a structure formed in large multinucleated osteoclasts from a ring of “circumferential” podosomes (Oikawa et al., 2012). Less frequently, filopodium-like protrusions that formed after a transient expansion of podosome-like protrusions were observed. Wang et al have also observed finger-like extensions (~ 2.5 μm) in RAW264.7 cultures, which they termed fusopods (Wang et al., 2015). In that investigation, fusopods were the predominant structures, although some cells reportedly fused at contacts between broad surfaces. Thus, both IL-4-mediated formation of MGCs and RANKL-mediated osteoclastogenesis *in vitro* are initiated by a protrusion. Furthermore, in both types of cells, the membrane protrusions originate from sites of podosome accumulation. However, despite these similarities, the majority of fusion-competent protrusions in osteoclasts form from the stationary sealing belt, which is not a feature of large MGCs. Instead, fusion-competent protrusions in MGCs are associated with highly dynamic peripheral podosomes (Figure 2A).

We demonstrated that macrophage fusion depends on F-actin and regulators of actin polymerization Cdc42 and its downstream effector WASp, inasmuch as the ability of macrophages to fuse was strongly impaired in the presence of cytochalasin B and in macrophages derived from WASp- and conditional Cdc42-deficient mice (Figures 7 and 8). In addition, inhibiting Arp2/3 activity also inhibited macrophage fusion. The decrease in fusion observed in cytochalasin B- and Arp2/3-treated as well as WASp- and Cdc42-deficient macrophages correlated with a lack of visible protrusions. Importantly, fusion of WASp- and Cdc42-deficient macrophages was reduced not only *in vitro* but also *in vivo*, suggesting that protrusion-mediated fusion is not an artifact observed exclusively in cultured cells. It is known that macrophages derived from Wiskott-Aldrich syndrome patients are defective in chemotaxis and phagocytosis (Thrasher, 2002). The finding that WASp is also essential for fusion extends its known roles in macrophages.

Our finding that the majority of fusion events in macrophages involved the leading edge of at least one cell (Table 1) and the observation of numerous thin protrusions at the site of contact between apposing cells (Videos 7 and 8) are consistent with filopodia formation at this location. Indeed, thin finger-like filopodia are found at the leading edge of migrating cells and are an extension of the branched lamellipodia network (Svitkina et al., 2003). However, these protrusions did not seem to initiate fusion. Although extension and retraction of filopodia persisted for a prolonged period, fusion occurred only after a wave of podosomes arrived and concentrated at the leading edge. At that point one protrusion initiated fusion. Therefore, it may be possible to distinguish two types of protrusions during macrophage fusion: *bona fide* filopodia as pre-fusion protrusions and a fusion-competent protrusion that assembled in a region enriched in podosomes. Fusion-competent protrusions appear to have a different morphology being wider and shorter than filopodia (Figure 2D). Although the molecular composition and organization of the F-actin network within the two types of protrusions remains to be defined, our data suggest that several types of actin-based protrusions that have different functions may be involved in macrophage fusion.

Previous studies have shown that Cdc42, WASp and Arp2/3 have an important role in the formation of podosomes in macrophages (Linder et al., 1999; Dovas et al., 2009; Linder et al., 2011). In line with these investigations, our results demonstrated fewer podosomes in macrophages derived from WASp^−/−^ and conditional Cdc42^−/−^ mice (Figures 6 and 7). In addition, the number of podosomes was significantly decreased in macrophages treated with Arp2/3 inhibitors (Figure 8). Although podosomes were enriched at the site of fusion and abolishing activity of critical podosomal proteins impaired fusion, the mechanistic link between fusion-competent protrusions and podosomes remains to be established. Podosomes are formed at the ventral side of the cell and typically protrude vertically into the substrate (Labernadie et al., 2010; Labernadie et al., 2014; Proag et al., 2015; Linder and Wiesner, 2016). Our current data suggest that the force driving the formation of fusion-competent protrusions may be delivered laterally toward the apposing cell in order to initiate fusion. It is possible that podosomes may directly or indirectly generate protrusive force during the fusion process. We speculate that when podosomes arrive at the leading edge they anchor the ventral actin network while allowing the lateral actin network to continue extending forward. Under these conditions, the cell would be unable to protrude along the entire leading edge and thus may focus the protrusive force to a limited region. Since the apposing macrophage presents an obstacle for the elongation of the protrusion, the protrusion may generate pushing force and penetrate into the adjacent cell. It is presently unclear why only certain macrophages display directional movement of podosomes and how it is associated with the formation of a fusion-competent protrusion.

Long fusion-competent protrusions that we occasionally observed in our experiments (Fig. 1C and 1D) are visually reminiscent of tunneling nanotubes (TNT). TNTs are thin long membranous tubes with diameters of 50-800 nm connecting two cells that have been reported in numerous cell types, including macrophages (Rustom et al., 2004; Kimura et al., 2012; Onfelt et al., 2006; Hanna et al., 2017). Formation of TNTs requires F-actin and, as recently shown in macrophages, depends on the activity of Rac1, Cdc42 and WASp (Hanna et al., 2017). Despite the general requirement for F-actin and the activators of actin polymerization, there seems to be clear distinctions between TNTs and fusion-competent protrusions. In particular, we observed that contact initiated by a long protrusion with a neighboring macrophage was invariably followed by fusion. In contrast, TNTs that also form by extending long protrusions remain stable structures that connect two cells. Furthermore, while TNTs have been reported to form in short 4-hour cultures of RAW/LR5 macrophages (Hanna et al., 2017), fusion of peritoneal macrophages begins 9 hours after addition of IL-4 (Faust et al., 2017). Finally, as revealed in our studies, short rather than long protrusions predominantly initiate macrophage fusion. Nevertheless, the requirement for F-actin and actin nucleation promoting factors of both TNTs and long fusion-competent protrusions is intriguing and suggests that two phenomena may be connected by a general mechanism.

The requirement for actin-based protrusions as initiators of cell-cell fusion seem to emerge as a unifying principle in several model systems, including fusion in osteoclasts (Oikawa et al., 2012) and fusion of muscle cells in flies, zebrafish and mice (Chen, 2011; Abmayr and Pavlath, 2012; Gruenbaum-Cohen et al., 2012; Aguilar et al., 2013). In *Drosophila*, muscle fibers are formed through rounds of fusion between a myotube (founder cell) and a fusion-competent myoblast (FCM). The fusion interface between these cells was found to contain an F-actin enrichment referred to as a “focus”, which develops in FCM and then invades the myoblast with one or several finger-like protrusions (Abmayr and Pavlath, 2012; Aguilar et al., 2013). Numerous actin regulatory proteins, including WASp, WIP, SCAR/WAVE and others have been found at the fusion site (Chen, 2011; Aguilar et al., 2013; Abmayr and Pavlath, 2012). Based on their invasiveness, size and the presence of the actin core with a surrounding ring of adhesive proteins, these structures were called “podosome-like structures” (PLS) (Sens et al., 2010). While the molecular composition of the fusion-competent protrusions in macrophages and myoblasts may differ, their formation nevertheless requires Cdc42, WASp and Arp2/3. Furthermore, although we did not observed stable F-actin foci in macrophages, fusion was initiated from sites where podosome clustered in large numbers (Figures 2A and 2B). Recent studies have shown that actin-based protrusions reconstituted together with adhesion molecules in *Drosophila* cells that normally do not undergo fusion, failed to recapitulate cell-cell fusion (Shilagardi et al., 2013). Expression of authentic fusion proteins in addition to cell-cell and cell-matrix adhesion molecules was necessary to induce fusion. These findings suggest that actin-based protrusions are insufficient on their own to promote cell-cell fusion. Whether force production alone is sufficient or the presence of fusion proteins is required for macrophage fusion is currently unknown. Further studies of actin dynamics may help define a link between actin-based protrusions and podosomes in macrophage fusion.

## Experimental Procedures

C57BL/6J, WASp^−/−^(B6.129S6-Was^tm1Sbs^/J) and Cdc42^loxP/loxP^ mice were purchased from The Jackson Laboratory (Bar Harbor, MA). LysMcre mice were a gift from Dr. James Lee. LifeAct mice (Riedl et al., 2010) were a gift from Dr. Janice Burkhardt and used with permission from Dr. Roland Wedlich-Söldner. The conditional Cdc42^loxP/loxP^ mice were generated by crossing Cdc42^loxP/loxP^ mice with LysMcre mice and screening for Cdc42 excision in myeloid leukocytes. All animals were given *ad libitum* access to food and water and maintained at 22 °C on a 12 light/dark cycle. Experiments were performed according to animal protocols approved by the Institutional Animal Care and Use Committees at Arizona State University and Mayo Clinic, Arizona and HHMI Janelia Reseach Campus.

### Macrophage isolation

Age and sex-matched mice were injected (I.P.) with 0.5 mL of a sterile 4% solution of Brewer’s thioglycollate (TG) (Sigma Aldrich, St. Louis, MO). All animals were humanely sacrificed 72 hours later and macrophages were isolated by lavage with an ice-cold solution of phosphate-buffered saline (PBS, pH 7.4) supplemented with 5 mM EDTA. The cells were collected into tubes pre-coated with BSA. Macrophages were counted with a Neubauer hemocytometer immediately thereafter.

### IL-4 induced macrophage fusion

Macrophage fusion was induced as previously described (Faust et al., 2017; Faust et al., 2018). Briefly, cells were applied to various surfaces at a concentration of 5×10^6^ cells/mL in DMEM/F12 supplemented with 15 mM HEPES, (Cellgro, Manassas, VA), 10% fetal bovine serum (FBS; Atlanta Biological, Flowery Branch, GA) and 1% antibiotics (Cellgro, Manassas, VA) and incubated in 5% CO_2_ at 37 °C for 30 min. Non-adherent cells were removed by washing the culture 3-5 times with Hank’s Balanced Salt Solution (HBSS; Cellgro, Manassas, VA) supplemented with 0.1% bovine serum albumin (BSA). HBSS was removed and the cells were incubated in culture medium for 2 hours. 10 ng/ml of IL-4 (Genscript, Piscataway, NJ) was applied to cultures until the respective time points. The fusion index (McNally and Anderson, 1995) was used to determine the extent of macrophage fusion.

### Phase contrast videomicroscopy

Macrophages were cultured on Permanox™ plastic (Thermo Scientific, Waltham, MA), Fluorodishes (World Precision Instruments, Sarasota, FL) or surfaces adsorbed with long-chain hydrocarbons as described previously (Faust et al., 2017). Dishes were transferred from the cell culture incubator to a stage-top incubator calibrated to maintain a humidified atmosphere of 5% CO_2_ in air at 37 °C. Phase contrast images were collected with a 20x or 40x objective every 30 s with an EVOS FL Auto (Thermo Scientific, Waltham, MA) and transferred to ImageJ to create movies.

### Lattice light sheet microscopy

The lattice light sheet microscope (LLSM) used in these experiments is housed in the Advanced Imaged Center at the Howard Hughes Medical Institute Janelia research campus. The system was configured and operated as previously described (Chen et al., 2014). Briefly, eGFP-LifeAct and/or mRFP-LifeAct peritoneal macrophages were applied to 5-mm cover glass surfaces adsorbed with long-chain hydrocarbons (Faust et al., 2017;Faust et al., 2018). IL-4 (10 ng/mL) was added and LLSM was conducted 8-10 hours thereafter. Samples were illuminated by lattice light sheet using 488 nm or 560 nm diode lasers (MPB Communications) through an excitation objective (Special Optics, 0.65 NA, and 3.74-mm WD). Fluorescent emission was collected by a detection objective (Nikon, CFI Apo LWD 25XW, 1.1 NA) and detected by a sCMOS camera (Hamamatsu Orca Flash 4.0 v2). Acquired data were deskewed as previously described (Chen et al., 2014), and deconvolved using an iterative Richardson-Lucy algorithm. Point-spread functions for deconvolution were experimentally measured using 200 nm TetraSpeck beads (Invitrogen) adhered to 5 mm glass coverslips for each excitation wavelength.

### Biomaterial implantation

Animal surgery was conducted according to AVMA guidelines by protocols approved by both Mayo Clinic and Arizona State University. Segments (1.5 × 0.5 cm) of sterile polychlorotrifluoroethylene (PCTFE) were implanted into the peritoneum of age and sex-matched mice as described (Jay et al., 2007). Animals were humanely sacrificed 14 days later and explants were analyzed for the presence of MGCs. Prior to explantation, 2 mL of PBS containing 5 mM EDTA was aseptically injected into the peritoneum and cells in the peritoneum were collected by lavage. The number of cells in the peritoneum at the time of explantation was determined by counting with a Neubauer hemocytometer. Experiments were conducted in duplicate on 3 independent days (i.e. 6 total mice per experiment).

### Immunofluorescence

At the indicated time point, specimens were fixed with 2% formaldehyde in PBS for 30 min at room temperature. Samples were permeabilized with 2% formaldehyde, 0.1% Triton X-100 in PBS for 30 minutes, then washed 3 times with PBS containing 1% BSA (PBS-BSA). Samples were incubated overnight at 4 °C with primary antibodies and 15 nM Alexa Fluor 488-conjugated phalloidin (Thermo Scientific, Waltham, MA). The specimens were washed 3 times with PBS-BSA and incubated with Alexa Fluor-conjugated secondary antibodies overnight at 4 °C. Nuclei were labeled with DAPI according to the manufacturer’s recommendation (Thermo Scientific, Waltham, MA). Samples were mounted in Prolong Diamond (Thermo Scientific, Waltham, MA) and imaged with a Leica SP5 and Leica SP8 laser scanning confocal microscopes housed in the W.M. Keck Bioimaging Facility at Arizona State University.

### Statistical analyses

Unless indicated otherwise results are shown as mean ± standard deviation from three independent experiments. Multiple comparisons were made via ANOVA followed by Tukey’s or Dunn’s post-test using GraphPad Instat software. Where applicable, means were compared by Student’s T-test. Data were considered significantly different if p < 0.05.

### Online supplemental material

Fig. S1 shows the distribution of eGFP-LifeAct and Alexa 568-conjugated phalloidin in fixed and permeabilized macrophages. Fig. S2 shows eGFP-LifeAct puncta in macrophages containing rings of vinculin. Video 1 is a time lapse video showing a short phase-dense protrusion that initiates macrophage fusion. Video 2 shows a long phase-dense protrusion that initiates macrophage fusion. Video 3 shows fusion between the leading edge and cell body. Video 4 shows fusion between the leading edge and lagging end. Video 5 shows fusion between leading edges. Video 6 shows fusion with eGFP-LifeAct macrophages. Video 7 is a maximum intensity isosurface render of eGFP-LifeAct showing actin-based protrusions during the fusion process. Video 8 is a maximum intensity isosurface render of eGFP-LifeAct showing stabilized and subsequently expanding protrusion between fusing macrophages. Video 9 shows asymmetric integration of eGFP-LifeAct and RFP-LifeAct signals in a mixed population of eGFP/mRFP-LifeAct macrophages. Video 10 shows the integration of eGFP- and mRFP-LifeAct with separated eGFP and mRFP channels. Video 11 shows deconstruction of mixed LifeAct signal creating the partition between macrophages.

## Acknowledgement

The authors wish to thank the members of the Ugarova laboratory and the ASU/Mayo Clinic Center for Metabolic and Vascular Biology for helpful discussions. We wish to thank Satya Khuon at HHMI Janelia Research Campus for help with sample preparation for LLSM. During the preparation of this work, James Faust was supported by a T32 Fellowship (5T32DK007569-28). The AIC at HHMI Janelia is jointly funded by Howard Hughes Medical Institute and the Gordon and Betty Moore Foundation. This work was supported in part by National Institutes of Health grant R01 HL 63199 to T.P. Ugarova. The authors declare no competing financial interests.

## Author contributions

J. J. Faust, A. Balabiyev and N. P. Podolnikova performed experiments and analyzed data. J. M. Heddleston and Teng-Leong Chew analyzed data, and provided tools and reagents. D. P. Baluch provided tools and reagents and assisted with the acquisition of phase contrast live cell imaging data. T.P. Ugarova and J. J. Faust conceived the project, designed research, analyzed the data and wrote the paper.

