## Supplemental figures for "An actin-based protrusion originating from a podosome-enriched region initiates macrophage fusion"

### Supplemental material

Faust et al,

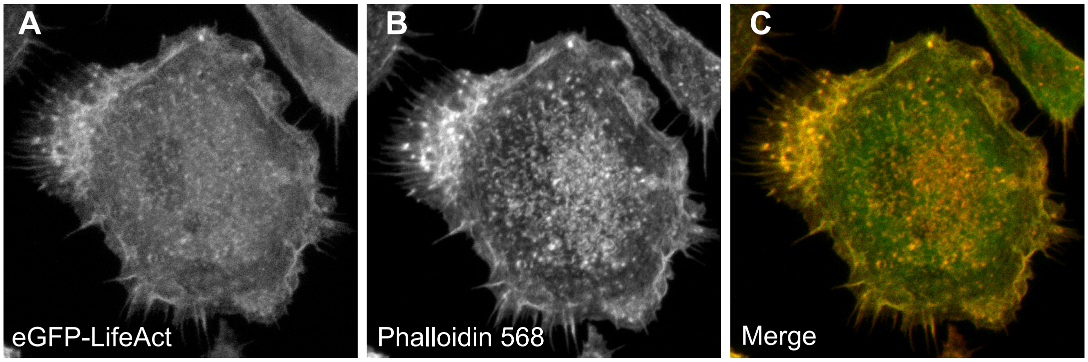


**Supplemental Figure 1: eGFP-LifeAct faithfully reports the distribution of F-actin in TG-elicited macrophages.** (A) The distribution of eGFP-LifeAct in a fixed and permeabilized macrophage 24 h after plating. (B) Alexa 568-conjugated phalloidin labeled structures. (C) The panel is an overlay of eGFP-LifeAct (green) and Alexa 568-Phalloidin (red). The majority of phalloidin-labeled structures appear to contain eGFP-LifeAct in fixed specimens.

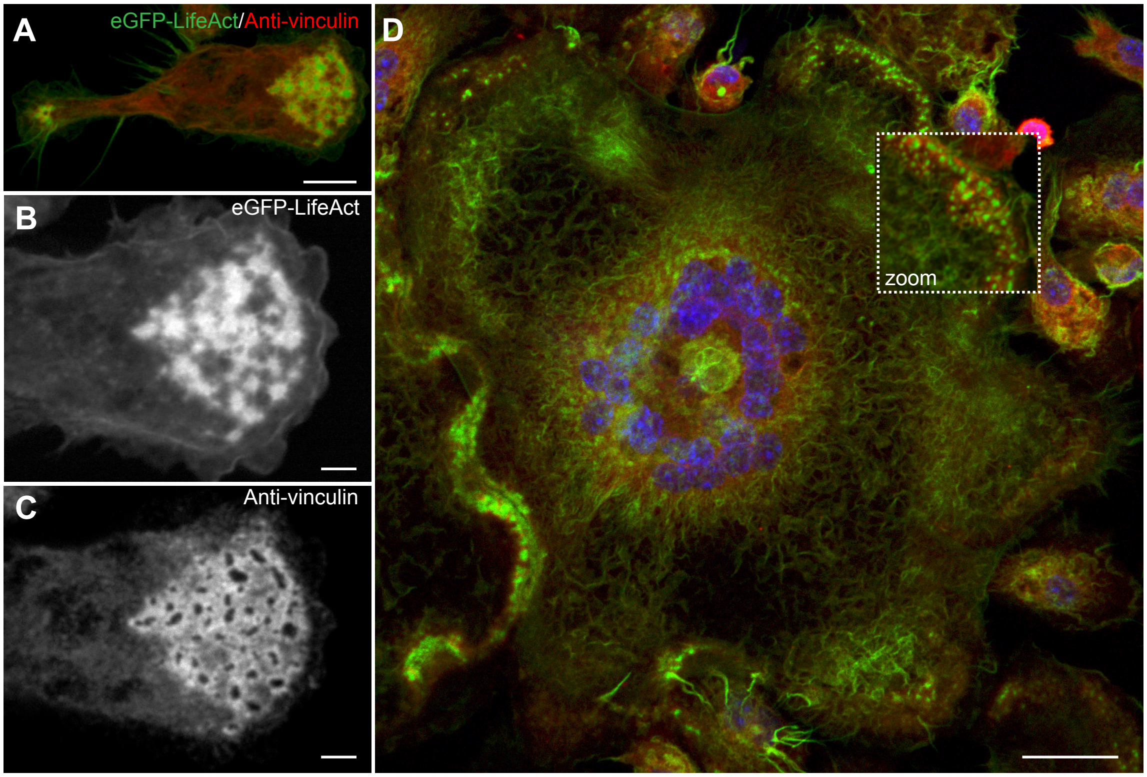


**Supplemental Figure 2: eGFP-LifeAct puncta in macrophages are podosomes.** (A) eGFP-LifeAct macrophage 48 h after the application of IL-4. Punctate eGFP-LifeAct structures (green) at the leading edge of a mononuclear macrophage contain anti-vinculin (red). The scale bar is 7.5 µm. (B) High magnification view of eGFP-LifeAct puncta shows the typical interconnected network with a core of eGFP-LifeAct and fine-fibrils radiating from a central point. The scale bar is 2.5 µm. (C) High magnification view of anti-vinculin enriched around a core devoid of anti-vinculin signal. The scale bar is 2.5 µm. (D) Low magnification view of a MGC shows similar podosomes at the leading edge. The scale bar is 25 µm.

**Video 1: A phase-dense protrusion initiates macrophage fusion.** Short phase-dense protrusions at the leading edge initiate fusion.

**Video 2: A long phase-dense protrusion initiates macrophage fusion.** Occasionally (~10%) long protrusions initiate fusion.

**Video 3: Fusion between the leading edge and cell body.**

**Video 4: Fusion between the leading edge and lagging end.**

**Video 5: Fusion between the leading edges**.

**Video 6: Lattice light sheet microscopy of fusing macrophages.** High spatiotemporal view of a MGC undergoing multiple fusion events is shown. A wave of eGFP-LifeAct puncta (white) emanate from the interior of the MGC and enrich at the cell periphery before fusion.

**Video 7: Maximum intensity isosurface render of eGFP-LifeAct.** Surface renders show actin-based protrusions during the fusion process.

**Video 8: Maximum intensity isosurface render of eGFP-LifeAct.** When there is adequate space between fusing macrophages fusion-competent protrusions can be observed. The protrusion appears to become stabilized and subsequently expand as fusion progresses.

**Video 9: Lattice light sheet microscopy of mixed populations of eGFP- and mRFP-LifeAct.** Integration of LifeAct signal appears to be asymmetric.

**Video 10: Lattice light sheet microscopy of mixed populations of eGFP- and mRFP-LifeAct.** eGFP and mRFP channels are separated to observe integration of eGFP- and mRFP-LifeAct.

**Video 11: Lattice light sheet microscopy of mixed populations of eGFP- and mRFP-LifeAct.** There appears to be deconstruction of mixed LifeAct signal that creates the partition between macrophages.
